# The impact of human brain geometry on the transport of an intrathecal tracer

**DOI:** 10.1101/2025.10.03.680193

**Authors:** Andreas Solheim, Geir Ringstad, Per Kristian Eide, Lars Magnus Valnes, Kent-Andre Mardal

## Abstract

**Background:** Intrathecal contrast-enhanced magnetic resonance imaging (MRI), utilizing the contrast agent gadobutrol as cerebrospinal fluid (CSF) tracer is emerging as a useful method to study glymphatic function in the human brain. A consistent finding with this technique is large inter-individual variability regarding tracer propagation. In this study, we outline an approach which predicts the distribution of tracer in the parenchyma based only on geometric information from brain tissue as captured by MRI, addressing one possible explanation for this variability.

**Methods:** Registrations are computed from pre-injection MRI, and used to map images at **24** hours after tracer injection to perform predictions of tracer enrichment in the parenchyma in other patients. We apply the method to a dataset of human brain MRI of **134** patients examined for different tentative diagnoses including idiopathic normal pressure hydrocephalus, spontaneous intracranial hypotension and idiopathic intracranial hypertension, as well as a group of reference patients.

**Results:** Tracer enrichment mapped between patients by image registration correlate strongly with actual observed enrichment. For patients in the reference group, the relative root mean squared error on our predictions is on average **26%** in the gray matter, and **15%** or less in other brain regions. Predictions are generally less reliable in the gray matter, and for patients with identified CSF leaks.

**Conclusion:** We show that predictions made from purely geometrical considerations correlate strongly with actual MRI tracer enrichment for patients with similar diagnoses, thus quantifying the role of geometry in tracer enrichment.

## 1 Introduction

The dual discoveries of the glymphatic [1] and meningeal lymphatic system [2, 3] have created a paradigm shift in our understanding of brain clearance mechanisms via the cerebrospinal fluid (CSF). Removal of metabolic waste from the brain via the CSF seems to be important for brain health and may be particularly tied to sleep [4–6]. Impaired clearance may cause accumulation of toxic proteins, and increased risk of dementias such as Alzheimer’s [7, 8] or Parkinson’s [9] disease. However, the mechanisms and efflux paths governing transport via the CSF in humans remain only partially understood. Current efforts in computational modeling have yet to fully explain the processes involved in glymphatic transport [10–17]. Recent efforts to predict solute transport in the brain by data-driven modeling are promising [18], however the low number of unique patient samples and the high-dimensional nature of current imaging modalities make such approaches challenging.

Lumbar intrathecal contrast-enhanced MRI, often denoted glymphatic MRI (gMRI), which includes repeated imaging over several days provides a unique perspective on the clearance of MRI contrast, as a surrogate for glymphatic function. In these studies, the intrathecal MRI contrast agent gadobutrol at a dose of 0.50 mmol, or less, has been used as a CSF tracer [19, 20]. This tracer is hydrophilic and contained extravascular when injected intrathecally, Accordingly, the molecule is expected to exit the brain via the CSF through the same pathways as toxic waste proteins which are not cleared primarily over the blood-brain barrier. Accordingly, by performing MRI at several time-points following injection, previous studies have shed light on the pathways of such molecules in the brain [19, 20]. Understanding the distribution of tracer in the brain may therefore be an important step towards better design and application of intrathecally administered drugs. This has been proposed as a possible pathway for treatment of a range of diseases in the central nervous system (CNS) [21–25]. However, studies of clearance following intrathecal tracer injection show significant individual variations within patient groups [26, 27]. Accordingly, examining the role of unique individual properties such as brain geometry may be a key part of untangling the properties of solute transport in the CNS.

In this work, we propose a method for predicting the distribution of MRI tracer in the brain following intrathecal injection using image registration. Image registration is an important tool in radiology, as it allows for more direct inter- and intra-subject comparisons of imaging data. Accordingly, many approaches have been proposed for brain imaging specifically [28–35], see Klein et al. [36] and Klein et al. [37] for a detailed comparison of some of these methods. Importantly, deformation fields computed by image registration provide an avenue for mapping quantities between patients. In our approach, deformation fields are computed by performing registration of *T*_1_-weighted MRI, obtained prior to tracer injection, from a dataset of patients to a target subject. The computed mappings are subsequently applied to images with tracer enrichment, recorded 24 hours following intrathecal administration. Accordingly, we outline an approach which predicts the distribution of tracer in the parenchyma based only on geometric information from brain tissue as captured by MRI. The same method was used in Solheim et al. [38] to map solutions from computational experiments between individuals, and build reduced order models. In this work, we build on this method to analyze data as obtained by *T*_1_-weighted MRI.

We apply our approach to a dataset of 134 individuals who underwent intrathecal contrast-enhanced MRI utilizing the contrast agent gadobutrol as a CSF tracer (gMRI); this was done for clinical reasons to assess evidence of CSF disorders. The diagnostic categories were defined after diagnostic work-up, including gMRI. The largest diagnosis group in the dataset are patients with idiopathic Normal Pressure Hydrocephalus (iNPH) a disease characterized by CSF flow disturbances and enlarged ventricles [39] with histopathological overlap towards Alzheimer’s disease [40]. Other patients are diagnosed with Spontaneous Intracranial Hypotension (SIH) identified by CSF leaks and Idiopathic Intracranial Hypertension (IIH). Patients without identified CSF disorders, or diagnosed with pineal or arachnoid cysts, are considered as a reference group. We perform a comprehensive voxel-wise analysis to understand the consequences of isolating the role of brain geometry in brain solute transport in the parenchyma.

## 2 Data and Methods

### 2.1 Data collection and preparation

#### 2.1.1 Patients and approvals

Participants in the study were all recruited while under clinical work-up for various CSF disorders, and imaged at the Department of Neurosurgery, University Hospital of Oslo-Rikshospitalet according to the protocol described in Ringstad et al. [20]. Study subjects underwent intrathecal gadobutrol injection as part of clinical assessment and were imaged using MRI before injection, in addition to several time-points following injection. Data examined in this study was collected in the period 2015 − 2019 and approved by the Regional Committee for Medical and Health Research Ethics (REK) of Health Region South-East, Norway (2015/96), the Institutional Review Board of Oslo University Hospital (2015/1868) and the National Medicines Agency (15/04932-7). The study was conducted following the ethical standards of the Declaration of Helsinki of 1975 (revised in 1983). Study participants were included after written and oral informed consent. No additional data was collected for this work. Parts of this dataset has been reported in previous works on glymphatic solute transport assessed with MRI [19, 20], but the role of brain geometry in tracer distribution has not been directly quantified before.

The present patient cohort contains 134 individuals, and we detail the characteristic data of subjects in the dataset in Table (1). The dataset contains a subset of individuals with identified CSF disorders, the largest of which are subjects diagnosed with iNPH, a neurodegenerative disease with symptoms like urinary incontinence, gait disturbance and dementia [39, 40]. While the iNPH diagnosis is made from a fuller picture of the condition of the patient, following the American-European guidelines [41], a characteristic of iNPH is enlarged cerebral ventricles associated with CSF flow disturbances [19, 42]. The dataset also contains two further groups of patients with identified CSF disorders: *N* = 15 patients are diagnosed with SIH with identified CSF leakage, and a further *N* = 16 with IIH. The remaining patients in the dataset are considered as a reference group (REF), and consists of patients who were not diagnosed with a CSF-related disorder following clinical work-up as well as patients diagnosed with pineal or arachnoid cysts. In studies of CSF clearance to blood the pineal and arachnoid cyst patients appear to display similar dynamics as the undiagnosed patients [26, 27] and accordingly these patients are included in the REF group. For the purpose of the present study, individuals in the REF group are considered as being an approximation of a healthy reference group.

**Table 1.** Characteristic data for patient groups in this study. Abbreviations: *iNPH*: idiopathic Normal Pressure Hydrocephalus, *IIH*: Idiopathic Intracranial Hypertension, *SIH*: Spontaneous Intracranial Hypotension, *REF*: Reference group

| Group | N | Sex (M/F) | Age | Height [cm] | Weight [kg] |
| --- | --- | --- | --- | --- | --- |
| iNPH | 49 | 32/17 | (61 ± 16) [24 – 80] | 175 ± 9 | 83 ± 18 |
| IIH | 16 | 2/14 | (34 ± 11) [19 – 57] | 166 ± 8 | 87 ± 16 |
| SIH | 15 | 4/11 | (48 ± 14) [24 – 72] | 171 ± 7 | 75 ± 20 |
| REF | 54 | 13/41 | (40 ± 14) [19 – 75] | 172 ± 8 | 82 ± 14 |
| **Total** | **134** | **51/83** | **(48 ± 18) [19 – 80]** | **172 ± 8** | **82 ± 17** |

#### 2.1.2 MRI protocol

Patients in this study underwent an MRI protocol, as introduced in Ringstad et al. [19, 20], aiming to illustrate brain-wide CSF transport and clearance by intrathecal injection of an MRI contrast agent. Following pre-contrast MRI, each patient had off-label gadobutrol intrathecally administered at a dose of 0.5 mmol (0.5 ml of 1.0 mmol/ml gadobutrol; Gadovist, Bayer Pharma AG) by an interventional neuroradiologist. After contrast injection, patients were imaged at multiple time-points during the first 24 hours, as well as at 48 hours and 4 weeks after injection to investigate any tracer retention. In this study, we only consider the post-contrast MRI performed at approximately 24 hours, which is when the concentration of this MRI tracer in the brain appears to peak. The pre-contrast image is used as a baseline for signal enhancement due to tracer enrichment. Due to practical considerations, post-contrast images could not be recorded at exactly the same time after intrathecal administration and the images we consider in this study were generally completed at *t* = (24 *±* 2) hours. In the interest of legibility, we refer to these images as being recorded at 24 h from this point on. For a more detailed clinical description of study execution, as well as details on the MRI procedure and scanner settings, we refer the reader to [20]. Longitudinal data was aligned using the FreeSurfer [43] software (version 6), and resampled to the FreeSurfer-standard 256 *×* 256 *×* 256 voxel grid with a uniform 1 mm resolution.

Brain regions of each patient were labeled automatically using FreeSurfer, based on 3D *T*_1_-weighted pre-injection MRI [43, 44]. We exclude the SAS and ventricles to evaluate tracer enrichment at 24 h only in the parenchyma. Due to the alignment of longitudinal data we apply the automated labels from the pre-injection image directly to the tracer enriched MRI.

### 2.2 Image registration

The fundamental idea of this work is to use image registration to compute intersubject mappings based on pre-injection *T*_1_-weighted MRI and subsequently apply these mappings to tracer enriched images. Given an image *I*_*i*_ and a target image *I*_*target*_, recorded before tracer injection, the goal of image registration is to compute 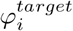, such that 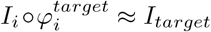. This mapping is based purely on information which can be derived from *T*_1_-weighted MRI, and therefore provides an avenue for translating quantities between individual subjects based only on information on brain geometry. As the contrast-enhanced images are performed in a relatively short time following injection, at the same time of day, we do not expect brain structure and geometry to change meaningfully in this time-frame. Accordingly, we use the mapping 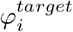, and apply it directly to the image recorded at 24 h post-injection, which we denote as 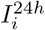.The resulting image 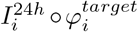 may therefore be seen as an estimate of 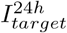 based solely on geometric information. In particular, if transport of the MRI tracer is entirely a result of geometric factors, then we expect 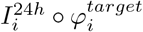 to approximate 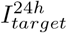, to the extent that 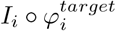 approximates *I*_*target*_.

In concrete terms, the method we propose in this work is the following: The dataset of pre-injection and tracer enriched MRI at 24 h may be seen as a set of *N* pairs 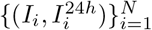. Given a target patient with pre-injection MRI *I*_*target*_, we compute the registration to this patient for each MRI in the dataset. We can thus compile a dataset of estimated tracer enrichment on the target patient brain geometry 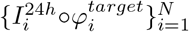. However, this does not account for cases where the physiological differences between patients are expected to be significant. The patient characteristics in Table (1) reflect that the present dataset is a highly heterogeneous group, with potentially meaningful differences in CSF flow and transport properties. In the following, we therefore split the estimated tracer enrichment at 24 h by patient diagnosis, as a proxy for capturing some of the geometric differences between patient groups. To generate estimated tracer signal predictions from different patient groups, we consider the median over each voxel in each patient group. We prefer the median over the average, in large part due to outliers in areas close to the SAS and ventricles, which are often insufficiently segmented in any single patient.

In the following, we perform image registration using the diffeomorphic SyN method from [34], implemented in the Advanced Normalization Tools (ANTS) tool-box (v.2.5.4). However, we emphasize that the method we propose is independent of the particular registration framework, as long as it allows for similar functionality. The performance of any given framework may depend importantly on the dataset in question, and accordingly we do not exclude the possibility that other frameworks yield similar or better results.

## 3 Results

### 3.1 Voxel-wise quantitative results

Using the approach outlined in Section (2.2) and illustrated in Figure (1), we generate group-wise predictions of tracer distribution at 24 h by taking the voxel-wise median of each patient group. In Figure (2) we show an example of the distribution profiles resulting from this approach, depicting a sagittal, axial and coronal slice showing the percentage signal change compared to the pre-injection MRI. The left column of this figure shows the data recorded in a clinical trial, while columns 2 − 5 are predictions generated using data from only the REF, iNPH, IIH or SIH groups respectively. In the right column, denoted as *All groups*, we show the prediction resulting from taking the voxel-wise median of all patients in the dataset which could be successfully registered to the target patient. The target patient in this particular example has no known CSF disorder, and may be considered as close to healthy, belonging to the REF group. We therefore particularly note the REF column in this figure, which presents an especially favorable comparison to the actual data in this case. This example also illustrates the variability of tracer distribution in different patient groups. Data from iNPH patients (column 3) appears to overestimate the signal increase in the gray matter, particularly noticeable in the anterior region of the sagittal slice. Meanwhile, data from the SIH patients (column 5) appears to underestimate the signal increase due to tracer injection compared to the actual data. In the following, we quantify these differences by performing voxel-wise statistical analysis.

**Fig. 1:**
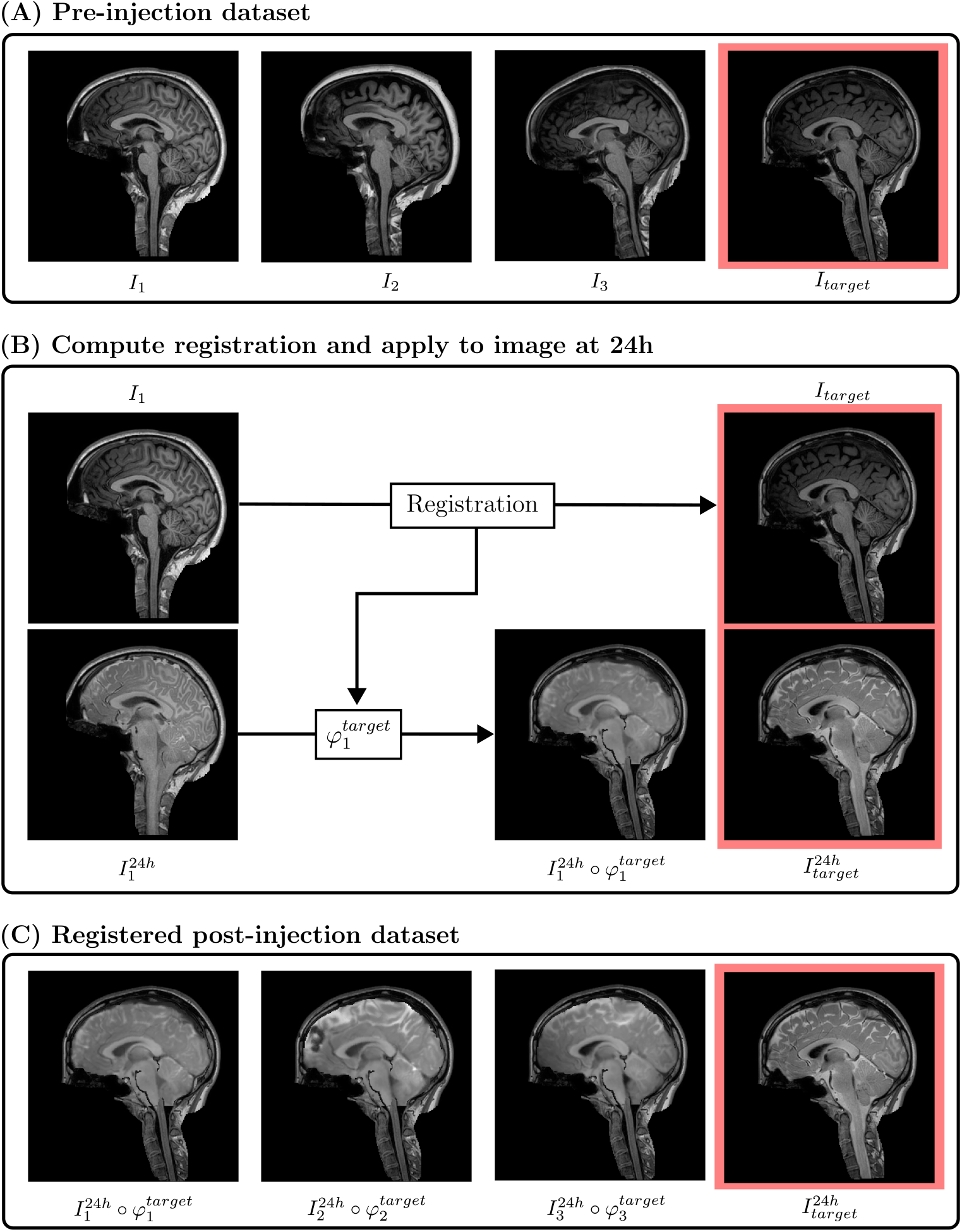
Stages of the image registration method. (A) Dataset of MRI prior to tracer injection. (B) For each subject in the dataset, we perform registration to the target patient based on the pre-injection MRI. The computed deformation field is applied to the tracer-enriched image at 24 h. (C) The method is applied to all subjects in the dataset, resulting in a dataset of post-injection registered tracer data.

**Fig. 2:**
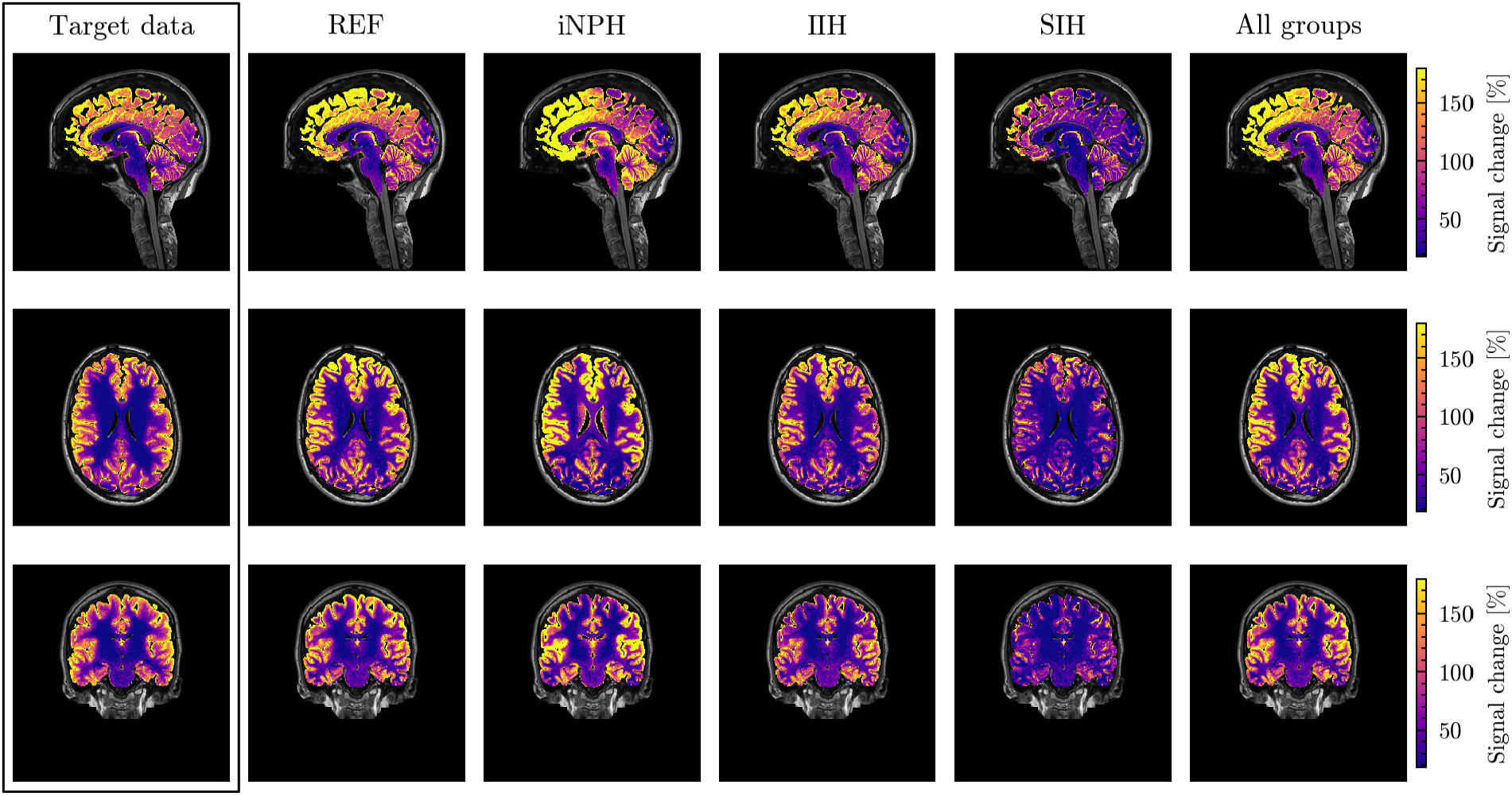
Clinical data compared to predictions of MRI signal increase 24 hours after intrathecal injection compared to pre-injection. The left-most column represents actual tracer enrichment observed in a clinical trial. Columns 2 − 5 represent voxel-wise median MRI signal in the four patient groups after registration to the geometry of interest. The right column represents the voxel-wise median of the entire dataset. The target brain is in the group of undiagnosed reference patients (REF).

To gain insight into the performance of geometry-based predictions on all patient groups in this dataset, we apply our method using 5 target subjects from each group. On a voxel-level we compute the signal-to-noise (SNR) ratio, relative root mean squared error (RMSE) and perform regression analysis of the target data against method predictions. To limit the effect of outlier values due to partial volume effects or other measurement deficiencies we exclude the 2% smallest and largest values when performing this analysis. The SNR is computed as the mean of these values divided by the standard deviation. Given target data **v** and prediction 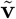 represented as vectors of *N* voxels, the relative RMSE is computed as 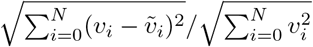. Linear regression is performed using the `linregress` function implemented in SciPy [45](1.15.2), with *p*-values estimated using the Wald Test with t-distribution of the test statistic using the null-hypothesis that the slope is 0. Each statistic is computed in the gray matter, white matter, cerebellum, limbic system and basal ganglia separately. Brain regions are identified from automatic segmentation of the target patient pre-injection MRI [44]. Figure (3) shows example correlation plots for the voxel-wise analysis we perform in the following. This figure corresponds to a voxel-wise view of the tracer profiles in Figure (2).

**Fig. 3:**
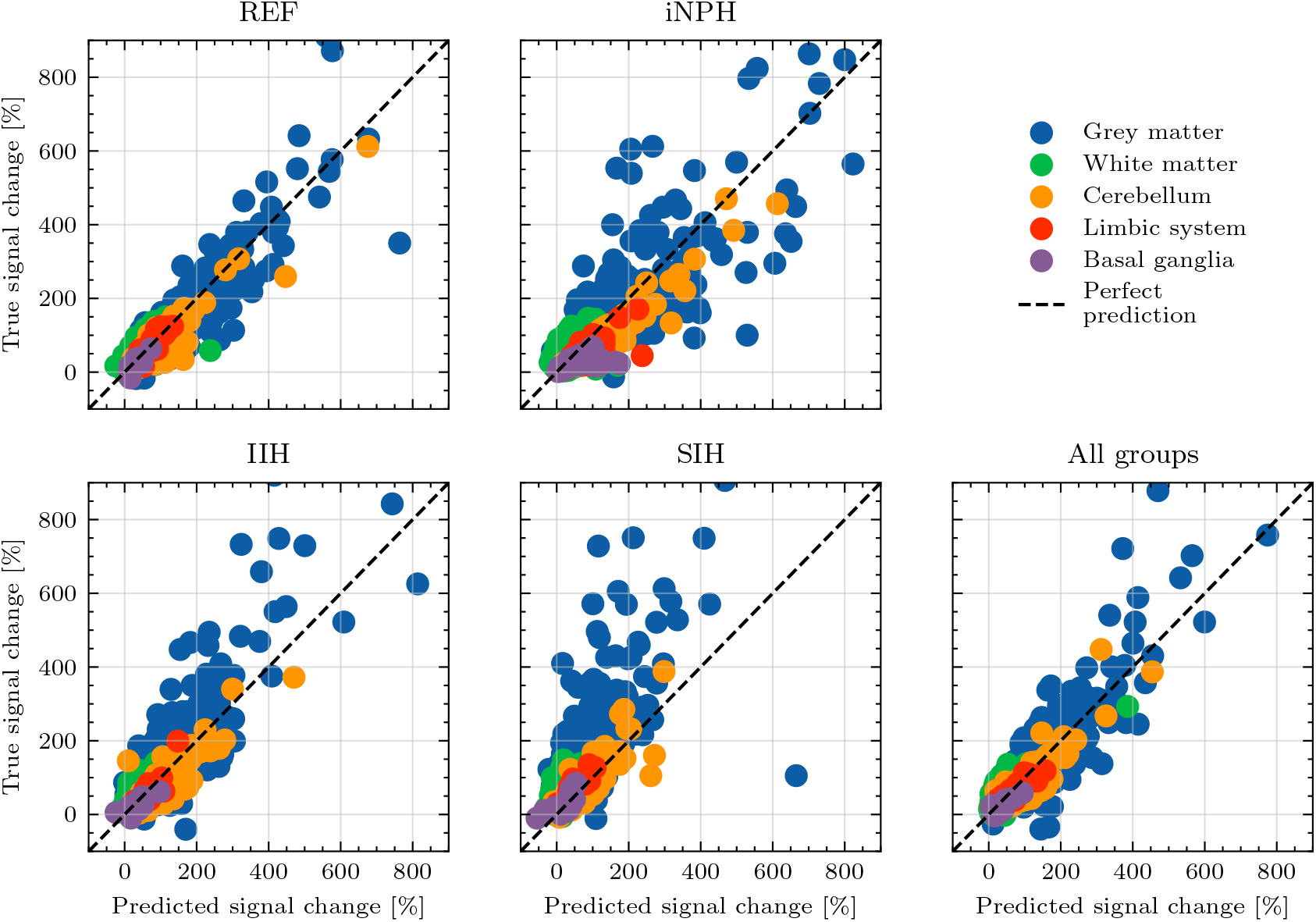
Predicted vs real signal increase in the parenchyma at 24 hours after intrathecal injection compared to pre-injection across all patient groups and five key brain regions. Each dot represents one voxel in the target brain MRI. In the interest of legibility we sample 0.01% of the voxels in each brain region uniformly at random. The target patient is REF and corresponds to Figure (3.1).

#### 3.1.1 Signal-to-noise ratio

In Table (2) we estimate the SNR for the target patient data in each of the 5 aforementioned brain regions. On the target data, shown in Table (2), SNR values generally lie in the range from 3 to 8. We find consistently lower values in the gray matter for all patient groups, while the white matter displays the lowest level of noise. The SNR in the cerebellum, limbic system and basal ganglia are similar across diagnosis groups, where we find values in the 5 − 6 range.

**Table 2.**
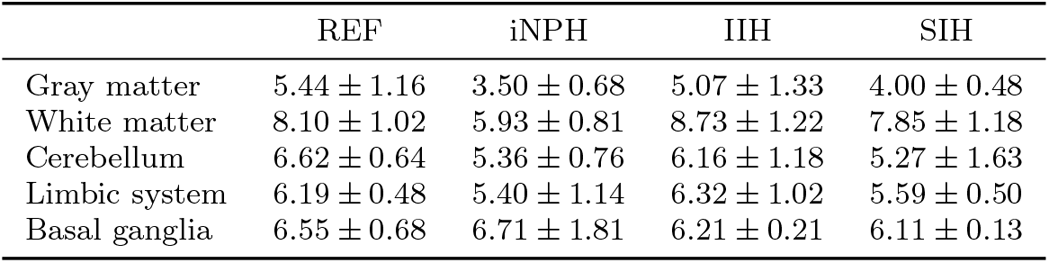
Average and standard deviation of the SNR of tracer enrichment at 24 h in each of the 5 target subjects in each patient group. The SNR is computed as the mean divided by the standard deviation in each brain region using values in the 2nd to 98th percentile range.

#### 3.1.2 Root mean squared error

To analyze the error of the predictions generated by image registration on a voxel-wise level, we display the average and standard deviation of the RMSE when our approach is applied to 5 subjects from each patient group, in Table (3). In particular, when performing registration to target subjects in the REF group, we notice that the RMSE is, on average, 26% or less in all regions when only using data from other REF patients. Errors are slightly larger when considering patients with a CSF diagnosis, but mostly remain less than 30% in regions outside the gray matter. Notably, the advantage gained by using data only within the same patient group seems to be diminished when the target patients do not belong to the REF group. When the target patients are IIH, seen in Table (3c), using data from REF patients appears to lead to better performance than using only IIH patients. Similarly, when the target patients are iNPH, as seen in Table (3b), data using data from iNPH, REF or IIH patients appear to lead to practically identical performance. This trend is however less clear when the target patients are diagnosed with SIH, as seen in Table (3d), where the error in the gray matter is typically significantly lower when using data from other SIH patients. Predictions generated for SIH target patients also show markedly larger average errors than all other patients groups. We also notice that the error tends to be subject to much greater variance when the method is applied to patients with CSF disorders. Comparing Table (3a) to Tables (3b–3d), the standard deviations tend to be meaningfully larger in the latter than the former, particularly in the gray matter.

**Table 3.**
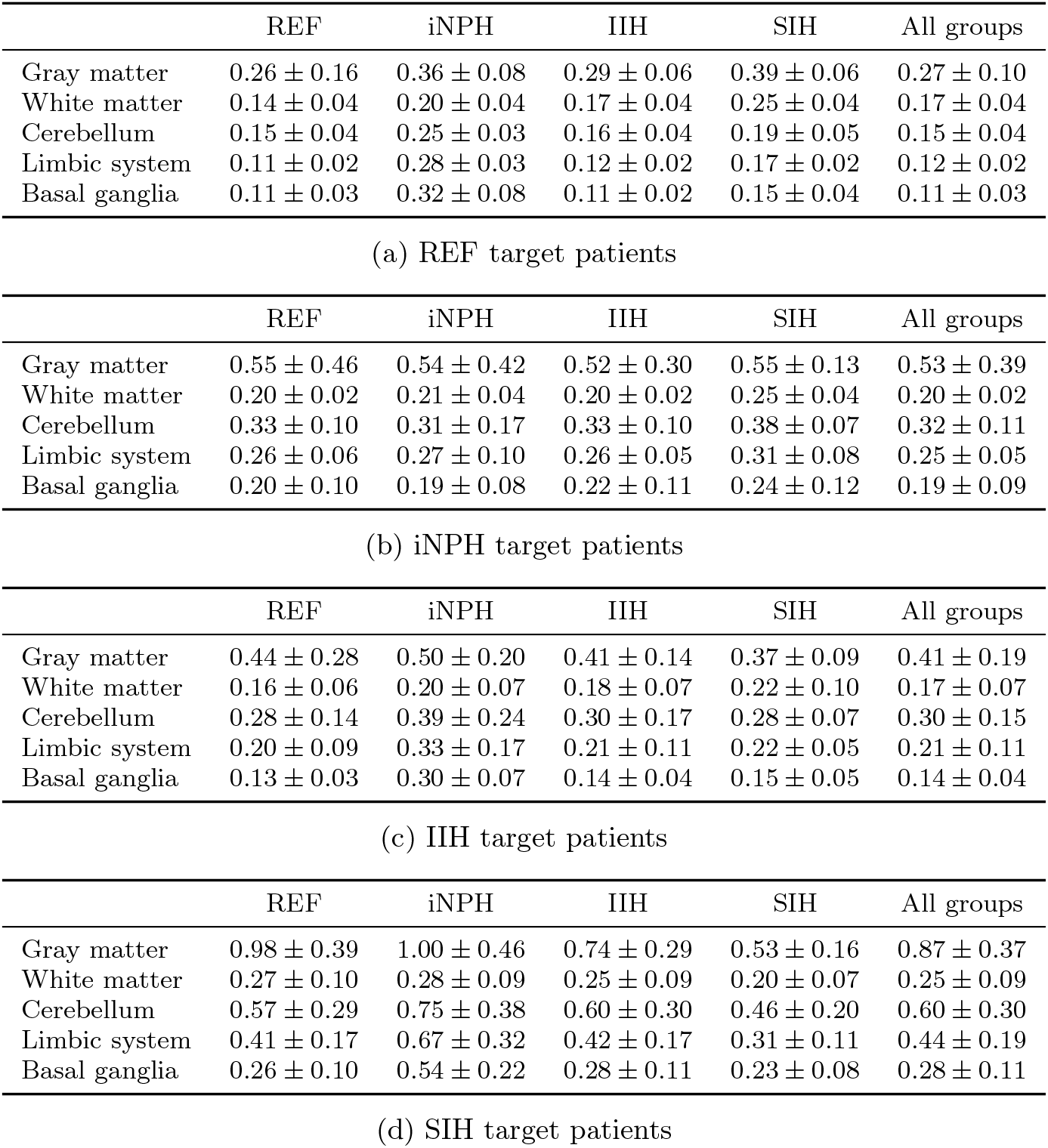
Average and standard deviation of the voxel-wise RMSE when performing registration to target subjects in the (a) REF, (b) iNPH, (c) IIH and (d) SIH groups. We examine data from 5 patients in each group.

#### 3.1.3 Regression analysis

In Table (4) we display computed Pearson’s *R*-values computed voxel-wise in the five aforementioned brain regions and using 5 target patients from each subject group. We display the average and standard deviation for each target patient group, and note that the *p*-values obtained when computing each *R*-value are in all cases smaller than the numerical truncation error due to the large number of voxels in each brain region. In Tables (4a – 4d), we find correlations which are generally stronger than 0.7, regardless of brain region. Correlations are typically weakest in the gray matter, and stronger in the basal ganglia and in the limbic system. We also bring attention to the correlations obtained in the basal ganglia when using iNPH patient data. It is particularly noticeable that when the target patients are not iNPH, i.e. Tables (4a), (4c) and (4d), the correlation is on average 0.33 or lower, which is much lower than in any other brain region. Meanwhile, when the target patients are themselves iNPH, we find correlations more in line with other brain regions, on average. As in Table (3), in many cases using only data from the REF group appears to lead to similar, or marginally stronger, correlations than when using patients with the same diagnosis as the target patients.

**Table 4.**
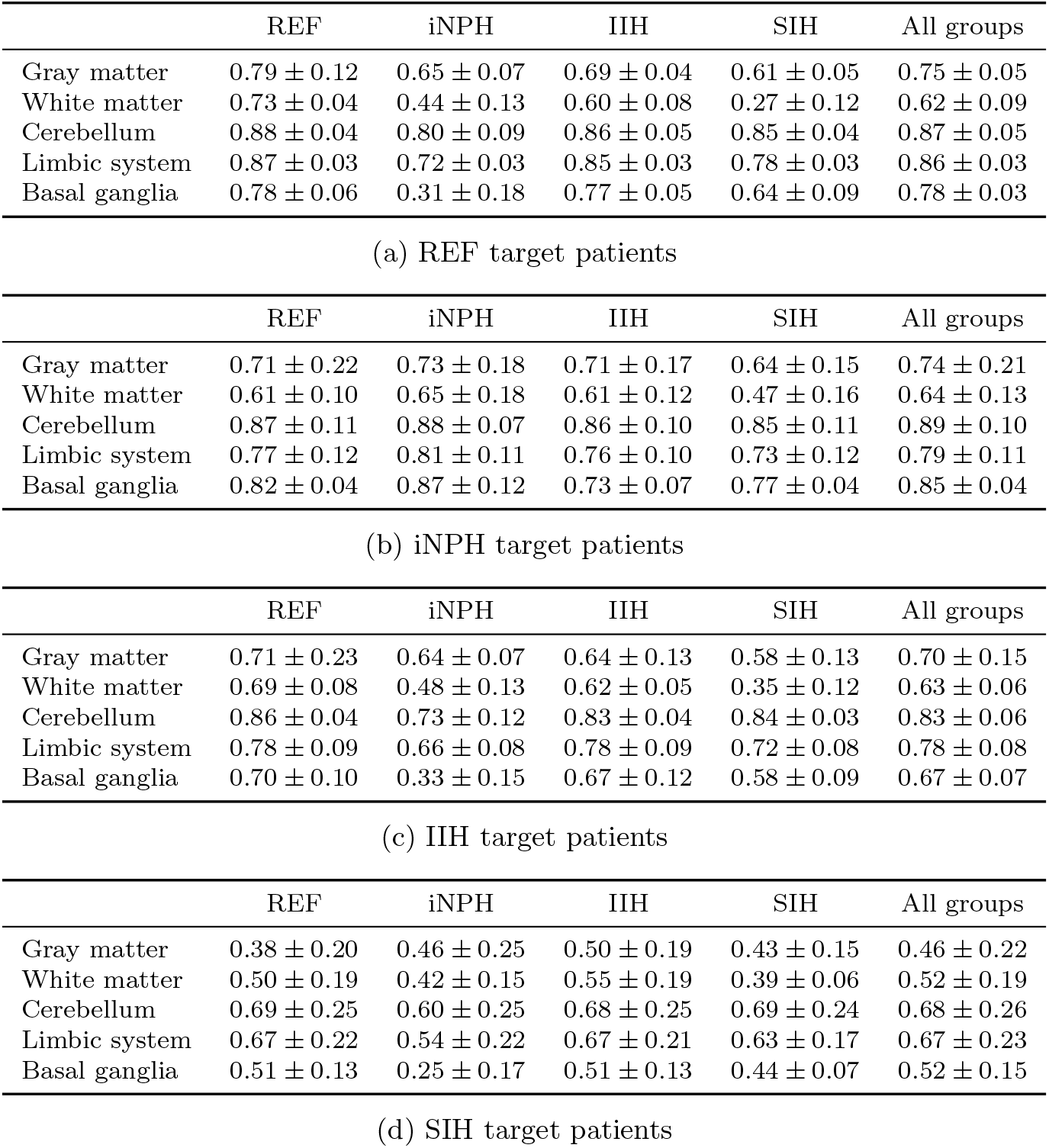
Average and standard deviation of the Pearson’s *R* correlation coefficient (*p <* 10^−16^ due to the large number of voxels in each region) when performing registration to target subjects in the (a) REF, (b) iNPH, (c) IIH and (d) SIH groups. We examine data from 5 patients in each group.

### 3.2 Qualitative examples

In Figure (4) we show examples of predictions generated from our image registration approach for four target patients, one in each patient group, compared to the true data recorded for that target patient. In each case, we use data from the patient group which results in the lowest relative error across brain regions in Table (3). Accordingly, for the REF and IIH target patients we use data only from REF patients, for the iNPH patient we use data from the entire dataset, while for the SIH target patient we only use data from other SIH patients. We particularly note the generally strong similarities between the predicted and actual tracer enrichment in Figure (4a). The tracer distribution pattern in the white matter of the axial and coronal slices are especially comparable. The main difference between the method prediction and the clinical data appears to be a slight overestimation of the tracer signal in the cortical gray matter, especially in the dorsal and anterior regions as seen in the sagittal and axial slices. For this particular target patient the relative error in the gray matter is 17%. Similar qualitative comments for the case where the target patient is IIH, seen in Figure (4c). In this case, however, unlike Figure (4a) the method appears to underestimate the tracer enrichment in the cortical gray matter. We also note more pronounced differences in the lateral directions in the axial and coronal slices. For the iNPH target patient in Figure (4b), we particularly note the tracer distribution in the cerebellum, which is strongly pronounced in the actual data measured on the target patient and not fully recovered by the prediction. Furthermore, we notice pronounced differences in tracer enrichment around the ventricles, visible in the axial slice. In the target data, the signal increase due to the MRI tracer is significantly larger than in the predicted profile. Figure (4b), also clearly shows the distinct morphological nature common in iNPH brains, where ventricles tend to become enlarged due to disturbances in CSF flow. Finally, in Figure (4d), we show data and method prediction for a target patient in the SIH group. As reflected in Table (3), patients in this group pose a particular challenge to this method, and we especially note the very low signal increase in the gray matter recorded in the clinical trial for this patient.

**Fig. 4:**
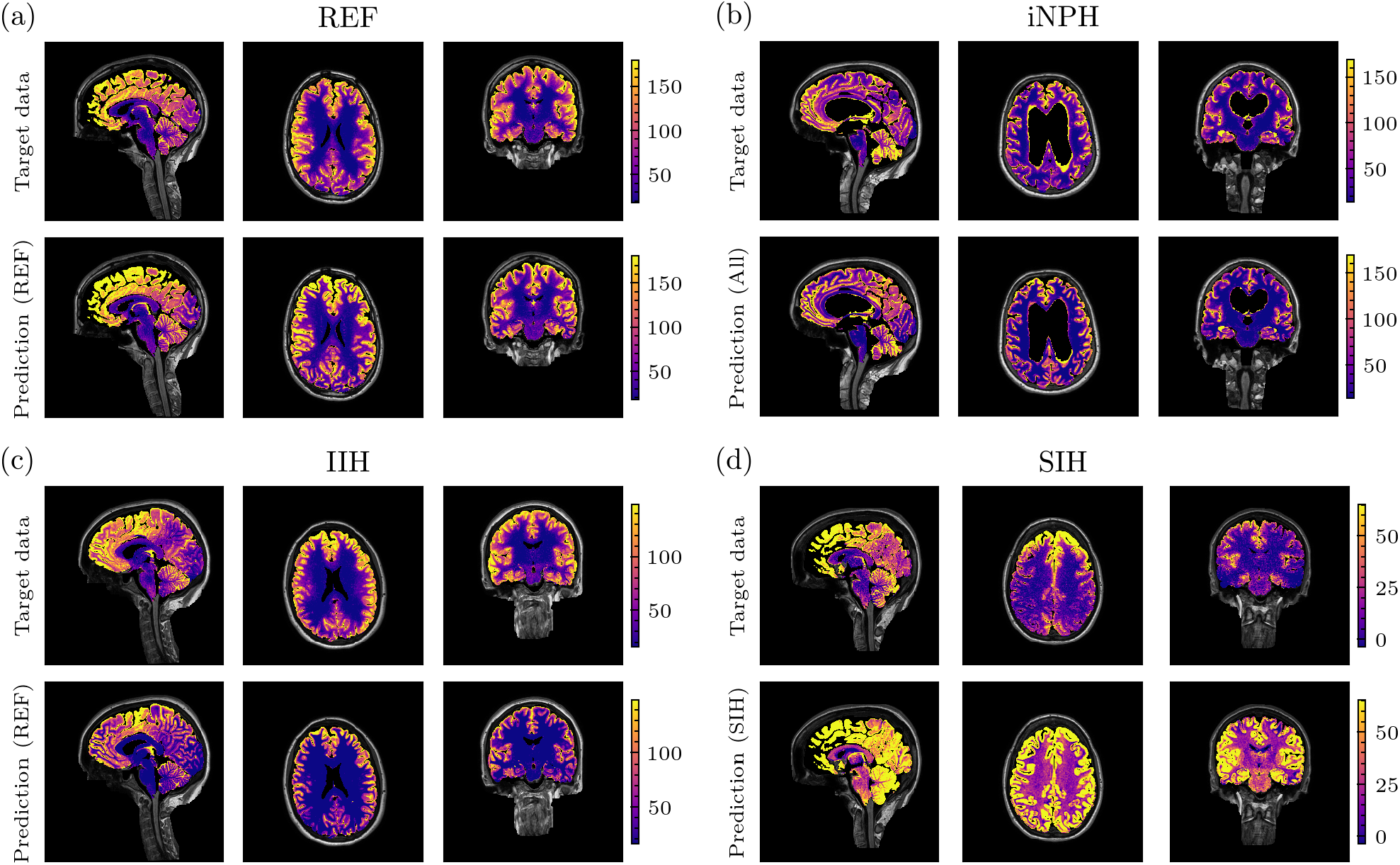
Best prediction generated using image registration compared to clinical data from the target patient for 4 individuals from the different patient groups. Color scale shows the percentage signal increase in MRI compared to the pre-injection image 24 hours following tracer injection.

## 4 Discussion

In this work, we have introduced a novel approach using image registration to predict the distribution of an MRI tracer in the brain parenchyma. *T*_1_-weighted MRI obtained prior to tracer injection is used to compute deformation fields from each patient in a dataset of individuals to a target patient. We then use these deformation fields to map the tracer distribution recorded at 24 hours for each patient to provide estimates on the target patient brain geometry. This allows us to isolate the role of brain geometry in transport of an intrathecal tracer. The method we present in this work therefore has the distinct advantage of being tied directly brain structure, and therefore more easily explainable than many data-driven methods. It is therefore a first step towards reliable, and verifiable, predictions of tracer distribution in the brain. Our approach is tested on patients diagnosed with iNPH, SIH and IIH, as well as a cohort of patients who were not diagnosed with a CSF disorder following clinical work-up and who are therefore considered to be approximately healthy (REF). Statistical analysis on a voxel-wise level was performed, to better understand the performance of such an approach.

### 4.1 Signal-to-noise ratio

The results in Table (2) show values for the SNR in the 3 − 8 range in all regions of the brain and all patient groups. This provides important context for the error analysis in Table (3). In particular, we note that for the REF target patient, when using data from other REF patients, the relative error generally reflects the SNR. In this case, we find that the prediction error is 11 − 26% on average, depending on brain region, which is what we expect for an SNR in the 4 − 6 range. The error on the REF patient predictions may therefore in large part be due to noise in target data. For patients with CSF disorder, this trend is not as clear. When the target patient is IIH, iNPH or SIH, we find relative errors from 40% up to 100% in the gray matter, which is greater than what the SNR computed for the gray matter, in Table (2), would indicate. In regions outside the gray matter, we find relative errors which are consistently lower, however, still higher than what the SNR indicates for these regions. For the three groups diagnosed with CSF disorders, we find SNR in the 5 − 8 range outside the gray matter, but the RMSE on our predictions are mostly higher than 20% for these patients. Accordingly, for these patients there are likely factors beyond noise which limit the capacity of purely geometrically based predictions.

### 4.2 Voxel-wise error

The results of this study illustrate the viability of a registration based approach as a basis for acting as a first order prediction of tracer enrichment in the parenchyma. Particularly for target patients in the REF group, in Table (3), the average relative RMSE is 26% on average in the gray matter and 15% or less in other regions when using only data from other REF patients. When the target patient is diagnosed with a CSF disorder, the relative error is generally higher, but our approach leads to mostly reliable results, with the SIH group as a notable exception. The results in this work are largely comparable with other studies like Vinje et al. [17], Valnes et al. [15] and Solheim et al. [38] which aim to model brain solute transport based on computational modeling. The approach used in [38] is also based on image registration, however, the method is applied to data from computational experiments and therefore subject to significantly less noise. In a computational setting, the authors find relative errors in the range 10% − 30% on an example problem where solution depends sharply on the cortical folds. We may also mention Rieff et al. [18], which applies a machine learning approach to the same dataset. The key advantage of this work is that we are able to make predictions in three dimensions, while [18] only applies their framework to two-dimensional slices. This enables us to retain key geometric features of the tracer distribution after 24 h, as seen in Figure (2) and especially Figure (4). While the results in this study are not sufficient to replace clinical trials, our approach may provide an avenue for initial analysis of patients scheduled to undergo intrathecal MRI tracer injection. Especially for patient without suspected CSF leaks, our approach provides a pathway towards a reliable and verifiable alternative to the full gMRI protocol. However, we emphasize the need for further work to establish more confident predictions, and a better understanding of methodological limitations.

It is notable that when using data for patients diagnosed with iNPH, IIH or SIH to predict the tracer distribution of patients in the same group, the error is generally significantly higher than when using REF patient data to predict data for other REF patients. This indicates that the diagnosis groups are in fact highly heterogeneous from a solute transport perspective, and possibly display greater individual variance than healthy individuals. A possible explanation is that the iNPH, SIH and IIH diagnoses are not made entirely on the basis of CSF flow dynamics. From a perspective of data-driven modeling, aiming to predict the distribution of tracer in the brain, it may therefore be advantageous to consider the patient diagnosis as a contributing input parameter, but not fully determinative of model behavior. Individual differences between patients, as seen in [26, 27], may also be due to factors not directly attributable to the brain. Physiological differences in CSF spaces, prior to the tracer ever reaching the parenchyma may play an important role, especially for SIH patients, which have identified CSF leaks. Additionally, as seen in Table (1), the number of patients differ importantly between groups. Results generated from REF patient data may lead to better results. in part due to the patient sample being larger, and therefore more representative.

We also emphasize that there are several additional error sources present in this approach, which are hard to distinguish from methodological errors. For instance, the image resolution is limited to 1 mm in each direction, which in many cases will be insufficient to fully distinguish brain tissue from CSF-space. This will result in partial volume effects, especially at the SAS boundary, where the MRI signal will be a combination of brain tissue and CSF-space. This may pose a challenge to the present approach as the MRI-signal in pure CSF, will be much stronger than in brain tissue due to the difference in volume fraction. Such effects may explain why the error is generally significantly larger in the gray matter in Table (3). We also note the post-injection MRI are not performed at exactly the same time, due to practical constraints. Images are mostly recorded within 1 − 2 hours of 24 hours after injection, however, the impact of this time differential is hard to quantify. Following the MRI at *∼* 24 h, patients were imaged again at *∼* 48 h and 4 weeks after injection, and it is therefore not possible to gauge the importance of a few hours in either direction. However, we note that pharmacokinetic modeling of CSF clearance to blood appears to be considerably shorter than 24 h [26, 27]. This leads us to assume that a few hours in either direction has only a minimal impact.

### 4.3 Regression analysis

The results in Table (4) show consistently very strong correlations between the MRI signal predicted using image registration and the true data measured in clinical trials, this is also reflected in the example correlation plot in Figure (3). In particular, regardless of the target patient diagnosis, the average correlation coefficient is 0.7 or higher in the vast majority of cases. The most noticeable exception to this trend are method predictions when using iNPH patient data in the basal ganglia. When the target patients are not iNPH we find considerably weaker correlations in this case. The reason for this behavior is likely a result of a combination of two factors: 1) The iNPH condition involves disturbances in CSF flow in the brain, particularly around the ventricles [39, 42], often causing ventriculomegaly. Accordingly, patients with this condition display a significant amount of ventricular tracer reflux [42], potentially causing different CSF flow patterns around the ventricles than other patient groups. This theory is further supported by the fact that the correlation appears to be 0.87 on average when considering iNPH patient data mapped onto iNPH target patients (Table (4b)). 2) Image registration is an optimization based approach, and accordingly brain-to-brain mappings will not necessarily exactly map points from a region in one brain to an equivalent region in another brain. We therefore expect such frameworks to typically fare worse on outlier brain morphology. In the particular case of patients diagnosed with iNPH, the ventricles will often be significantly enlarged compared to other patient groups, and these patients are therefore particularly ill-suited for image registration. A practical consequence of insufficient registration in this case may be that voxels which are located in the ventricles of the iNPH patients are mapped to voxels in areas inside the parenchyma, but close to, the ventricles in the relevant target brain. This may lead to a signal prediction which is too large in ventricle-adjacent areas.

From a perspective of data-driven modeling of tracer transport in the brain, the present approach is possibly most efficiently thought of as a useful pre-processing step for further statistical analysis. The main challenge for data-driven modeling of this dataset is the inherent challenge of dimensionality involved in this problem: The number of available patients is in the order of hundreds, while the number of voxels in each three-dimensional image is on the order of tens of millions. A major advantage of the approach we propose in this work is that each quantity in the dataset is mapped to a common geometry, essentially accomplishing a significant part of building an efficient embedding space, which is a key part of most data-driven methods. Establishing a method for controlling and isolating some of the effect of individual brain variations can be helpful in allowing more efficient subsequent statistical modeling.

### 4.4 Qualitative examples

The key insight offered by the registration-based approach presented in this work is the ability to leverage key geometric properties in the target patient brain morphology when making predictions. This is especially well illustrated by Figures (4a) and (4c), where the predicted profiles appear to recover the tracer distribution in the target data to a significant degree. We note the large signal increase predicted by our method near the cortical gray matter, similar to the target data, with an apparent decreasing gradient towards inner brain regions. The coronal slice in Figure (4a) is a particularly favorable example. Similar observations can be made for the iNPH target brain case, shown in Figure (4b), but we notice particular differences close to the ventricles in this case. In particular, the MRI signal appears to be especially pronounced in the clinical measurements around the outside of the left and right ventricles, as seen in the axial slice of this figure. This may be due to ventricular reflux or edema, which is common in the ventricles of iNPH patients, but could also be explained by erroneous segmentation. We particularly notice an area of large signal increase in the posterior part of the right lateral ventricle, visible in the axial slice of the target data in Figure (4b), which is not as pronounced in the predicted profile. In addition to segmentation errors, we may intuitively expect larger registration errors close to the ventricles due to their extreme morphology. The comparison of the brain geometry in Figure (4b) to Figures (4a), (4c) and (4d) is striking in this regard.

Meanwhile, the case which appears to be the most challenging for our current approach are patients diagnosed with SIH. This is quantified in Table (3), but Figure (4d) illustrates the issue with this patient group. It appears that for SIH patients, individual variations in the amount of tracer which reaches the parenchyma, or is cleared to blood, largely outweigh individual geometric variations. For this patient group, it would likely be critical to take advantage of additional information on the total tracer amount which is expected to be present in the brain.

### 4.5 Limitations

This work considers only tracer enrichment in the parenchyma at one point in time. The present approach can be readily extended to also include CSF spaces and other time-points. Extending the analysis to areas where signal from CSF dominates would likely also require registration of *T*_2_-weighted images or other CSF-specific imaging modalities to produce accurate results. MRI of tracer prior to 24 hours tend to show tracer enrichment primarily in CSF spaces, and accordingly these two further extensions are likely inextricably linked.

Additionally, we consider MRI performed at approximately 24 h following intrathecal injection, however the precise time of this measurement may vary by 1 − 2 hours. Certain patient groups also contain 16 or fewer individuals, limiting our understanding of larger cohort behavior. However, the main limitation in this regard is the lack of true reference patients which are fully healthy. Results in Tables (3) and (4), in particular, show that the present REF group is especially useful for making estimations of MRI tracer signal even when the target patient is diagnosed with a CSF disorder, but not vice versa. This indicates that healthy reference patients may be a valuable resource to better predict and understand the brain solute dynamics also of patients who are not themselves healthy.

## 5 Conclusion

In this work, we have demonstrated an approach based on an image registration framework for predicting the distribution of an MRI tracer in the parenchyma 24 hours after intrathecal injection, based only on geometric information and knowledge of patient diagnosis. We find that predictions based on geometric information correlate with clinical data in all patient groups and in most brain regions, particularly when considering data from the REF cohort. This method is therefore a promising way of isolating some of the individual variance involved in brain solute transport and may act as a starting point for further statistical modeling. By focusing on brain structure when making predictions of tracer distribution, we open a new pathway towards more reliable and verifiable methods for predicting tracer transport in the brain. While the present study is limited to tracer distribution at one time-point and only in the parenchyma, our method can be extended to analyze tracer distribution in CSF spaces, and in time.

## Declarations

## Acknowledgments

Computational experiments and data processing were performed on resources provided by Sigma2 — the National Infrastructure for High-Performance Computing and Data Storage in Norway, under grant grant NN9279K.

## Ethics and approvals

Parts of the data presented in this has been presented in previous works on MRI-based assessment of human glymphatic function conducted at the University Hospital of Oslo [19, 20] in the years 2015 − 2016. Collection of data analyzed for this study was approved by the Regional Committee for Medical and Health Research Ethics (REK) of Health Region South-East, Norway (2015/96), the Institutional Review Board of Oslo University Hospital (2015/1868) and the National Medicines Agency (15/04932-7), and conducted following the ethical standards of the Declaration of Helsinki of 1975 (revised in 1983). Study participants were included after written and oral informed consent. No new data was collected for the present work.

## Availability of data and materials

The dataset used in this study is not made available due to patient privacy concerns

## Authors’ contribution

ARS and KAM developed the method and designed the study. PKE and GR performed the gMRI protocol and obtained MRI images. LMV performed data processing and curation. ARS implemented the method, performed data analysis, designed the figures and wrote the initial manuscript. All authors contributed to the editing of the manuscript and figures. All authors read and approved the final manuscript.

